## Supplementary figures and images for "Uterus-specific transcriptional regulation underlies eggshell pigment production in Japanese quail"

### S1 Fig

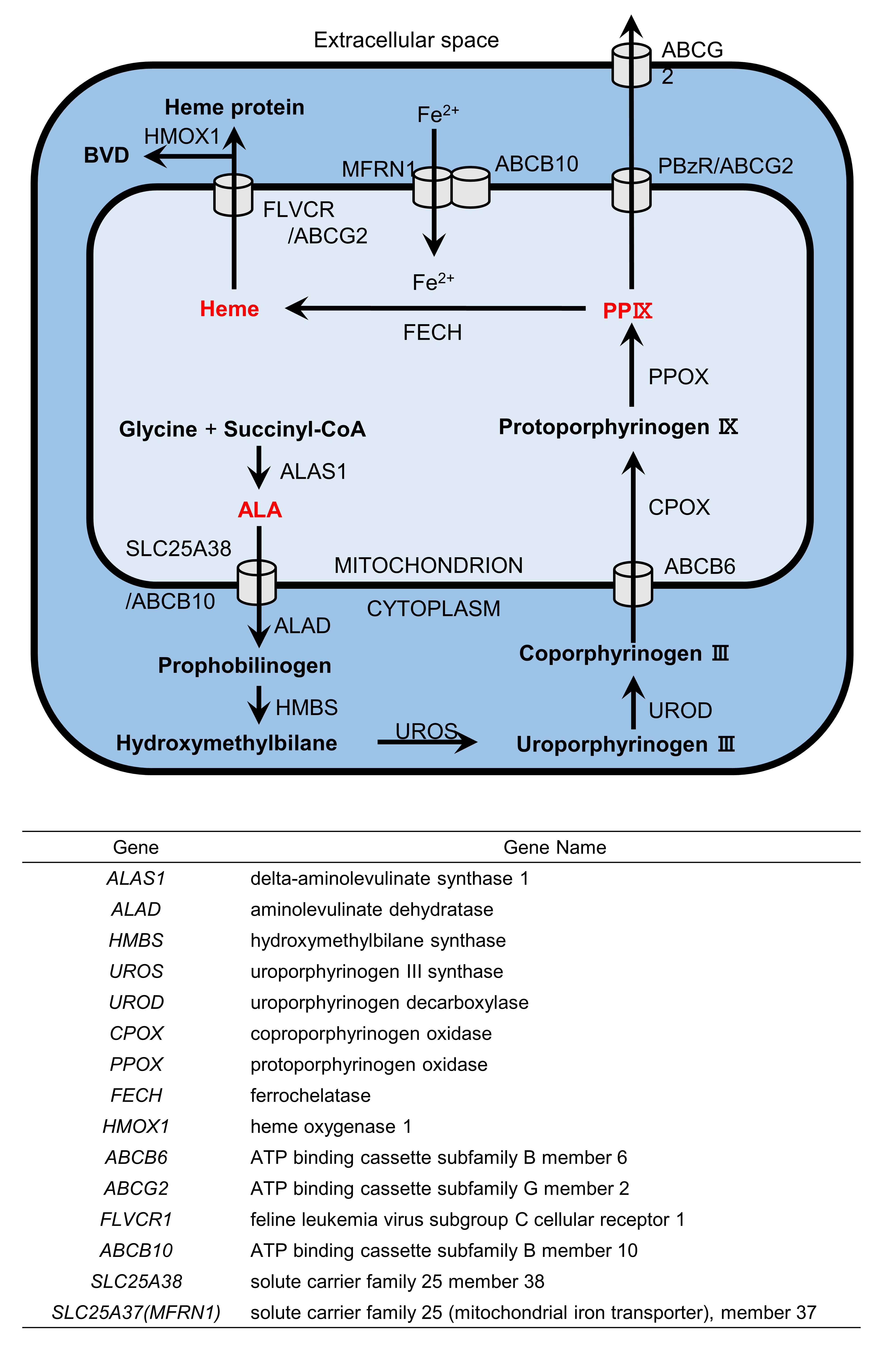

### S2 Fig

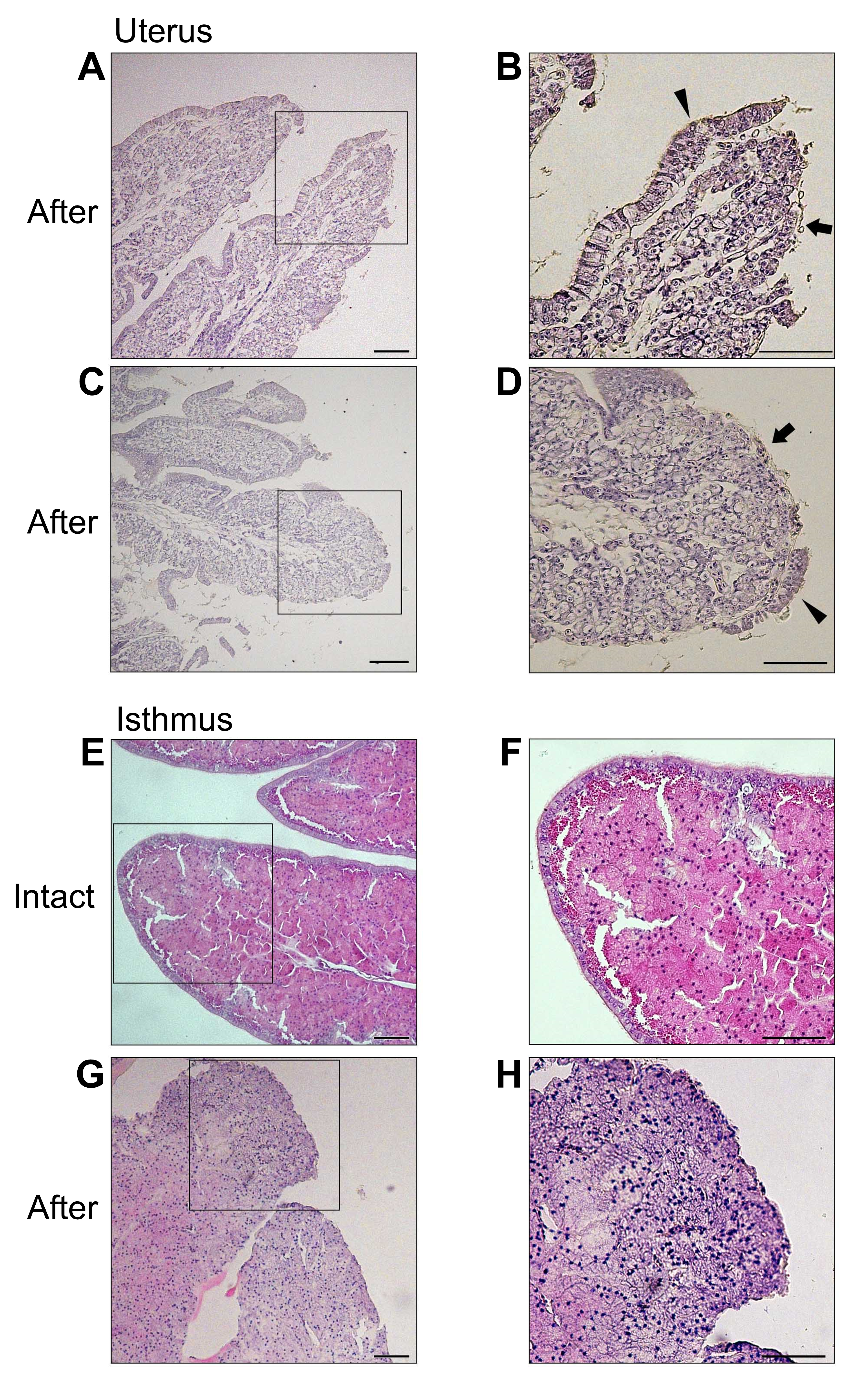

### S3 Fig

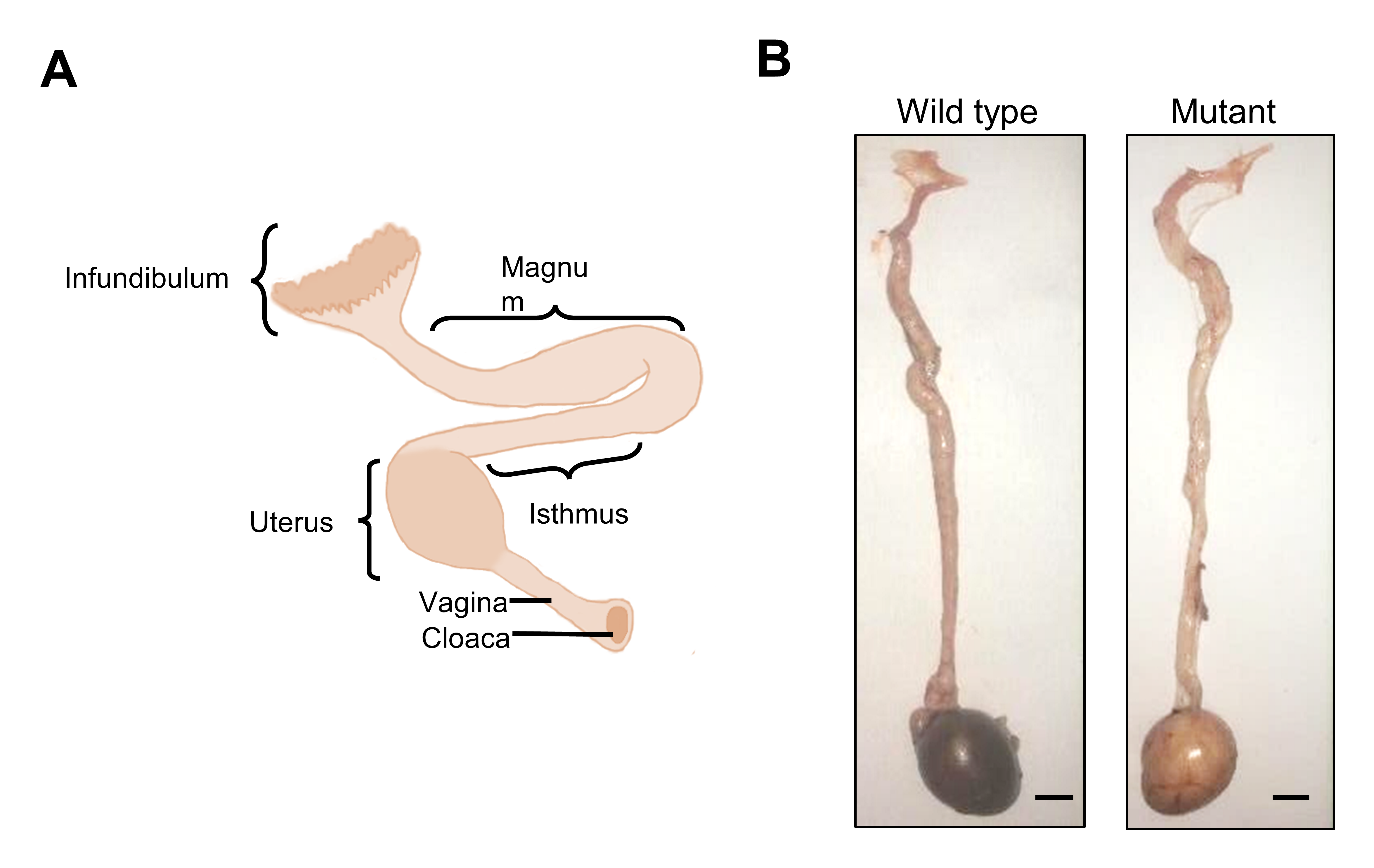

### S4 Fig

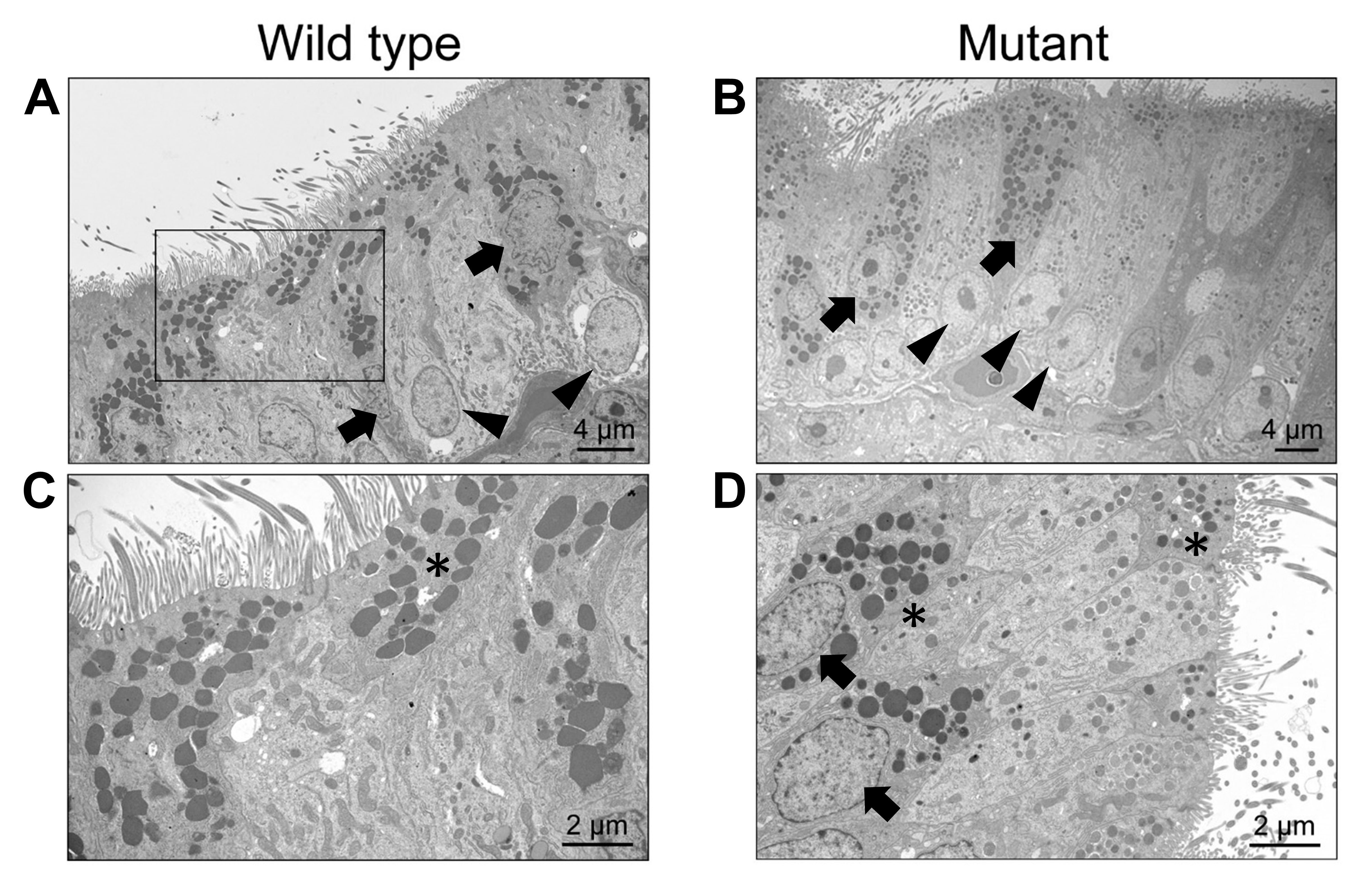

### S5 Fig

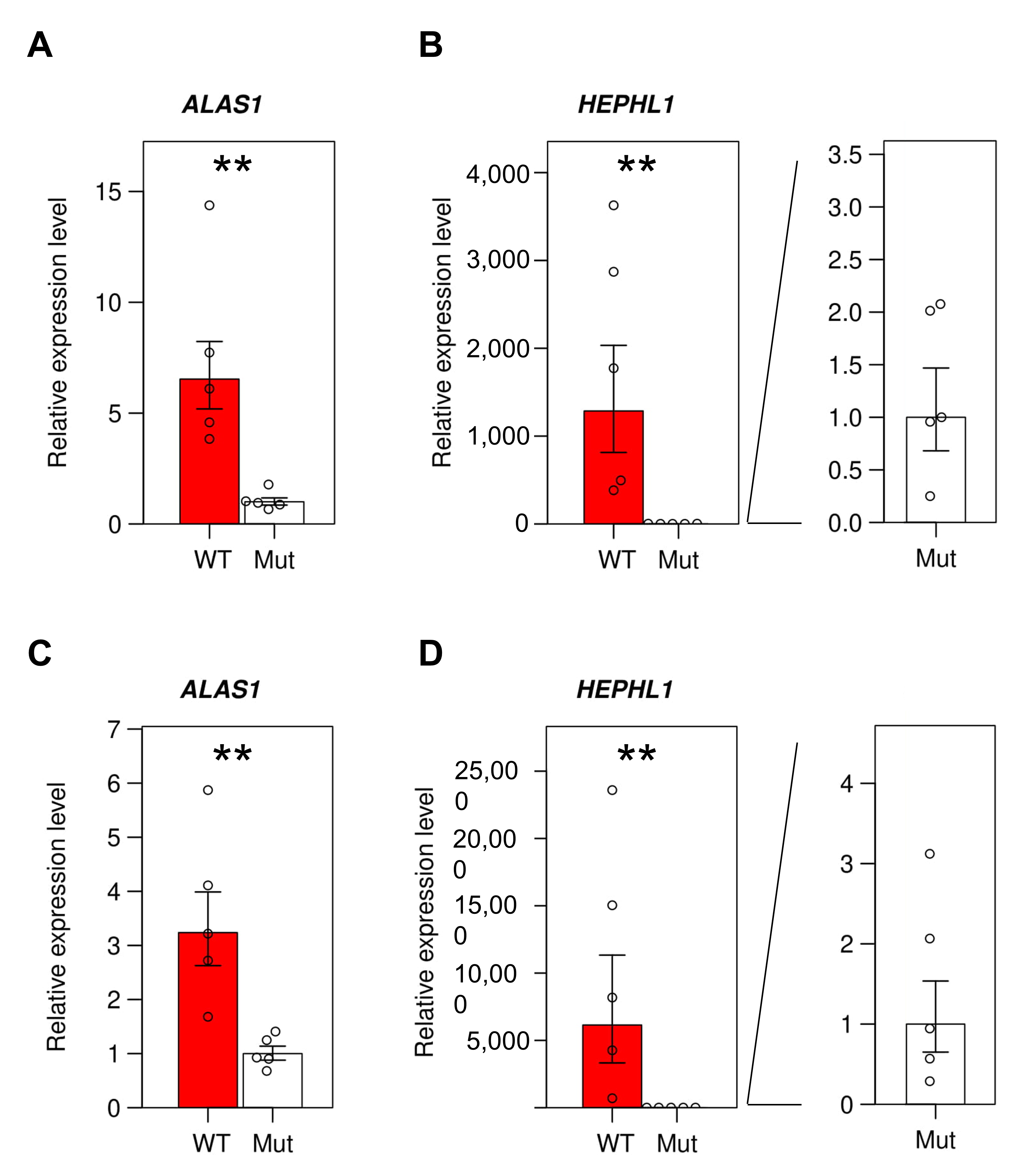

### S6 Fig

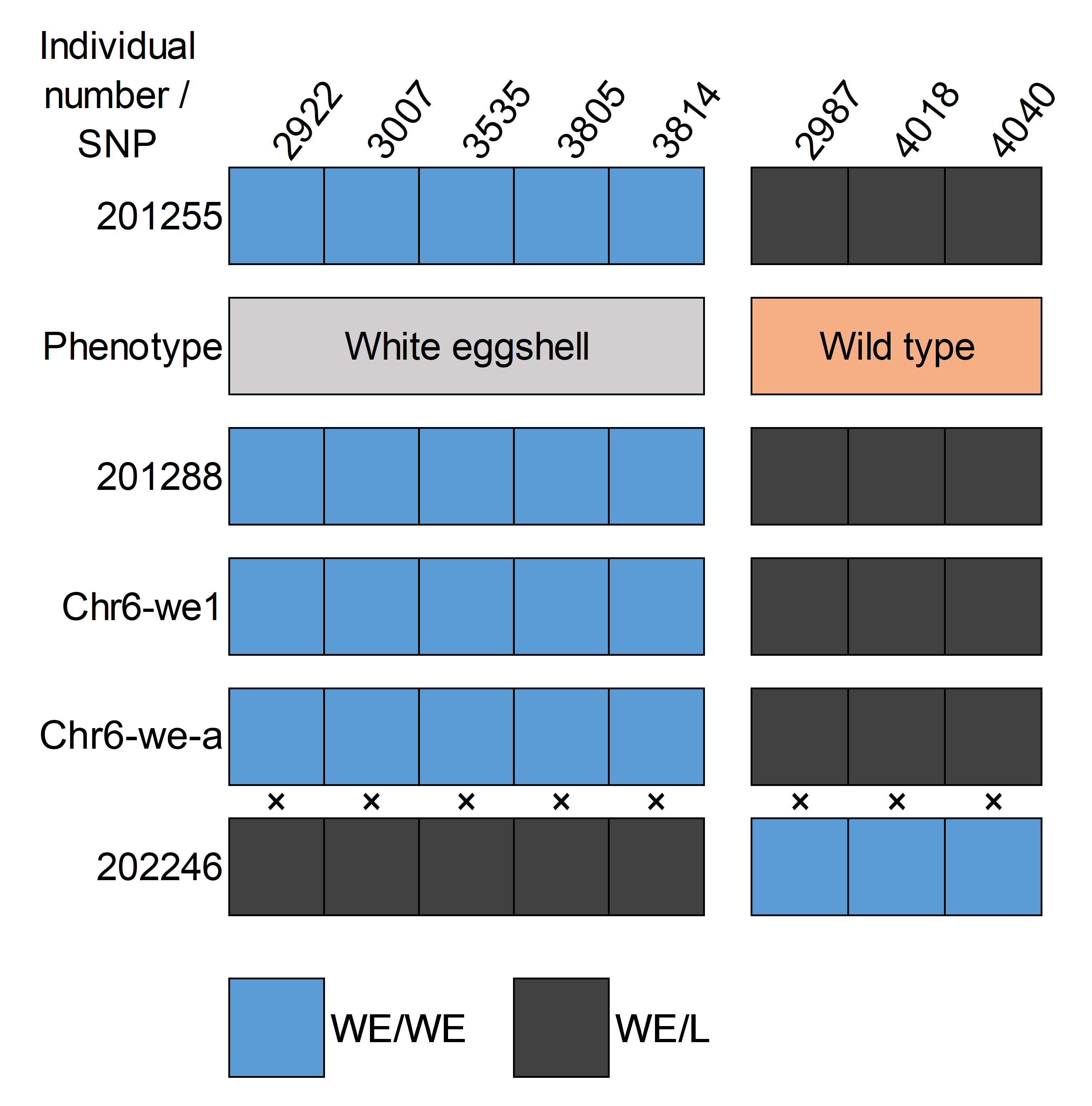

### S7 Fig

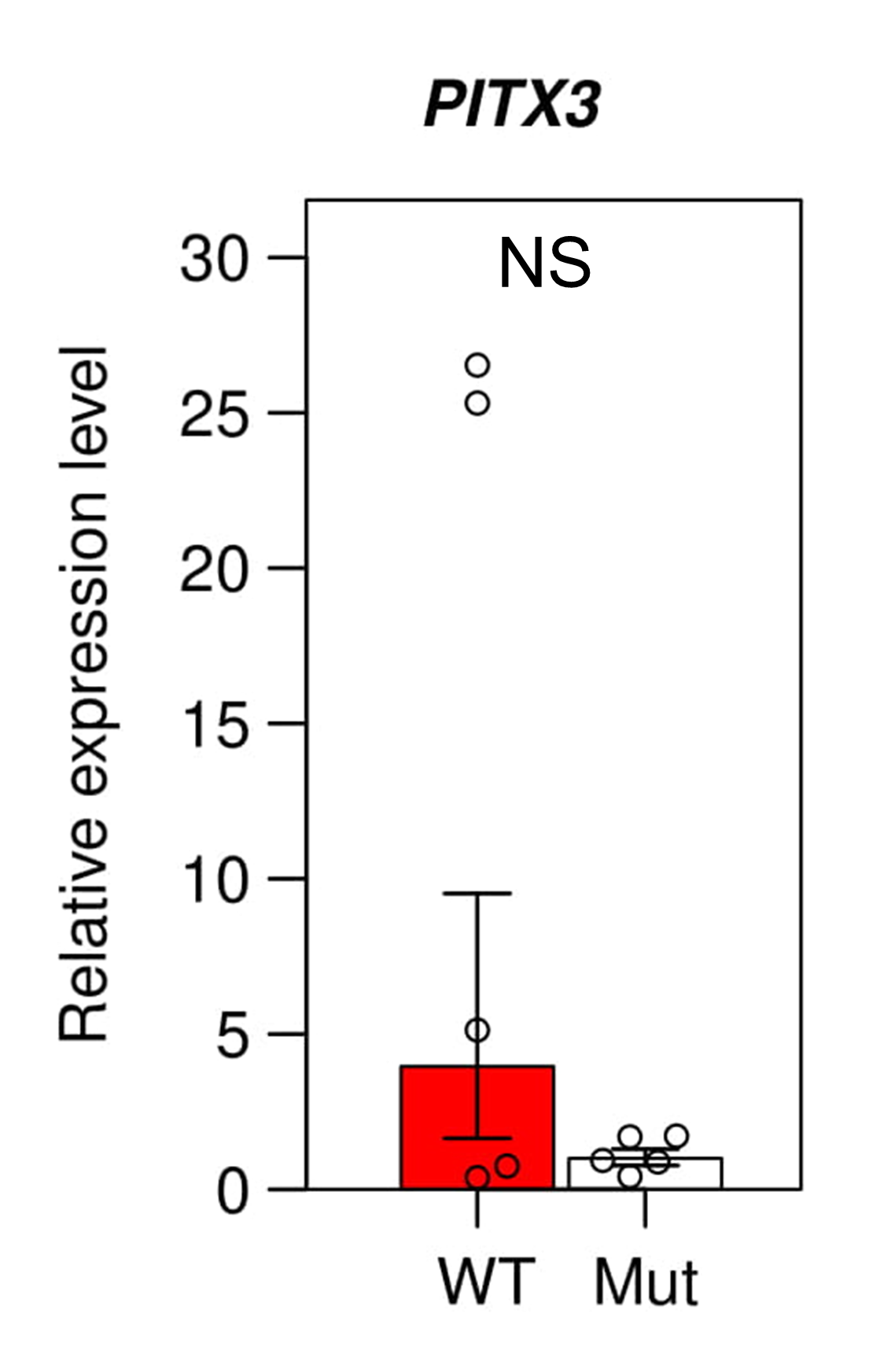
