## Supplementary material for "Uterus-specific transcriptional regulation underlies eggshell pigment production in Japanese quail": S2 Data

**Sequence of *ALAS1* cDNA used for *in situ* hybridization (nt): (1,298 bp)**

5′- CATACCTACAGAGTGTTCAAAACGGTGAACCGAAAGGCACAGATCTTTCCCATGGCAGATGACTACTCTGACTCCCTGATCACCAAGAAGGAGGTGTCTGTCTGGTGCAGCAATGATTACCTGGGCATGAGTCGTCACCCTCGTGTGTGCGGAGCAGTGATGGATACACTGAAACAACATGGTGCTGGAGCAGGAGGCACAAGAAATATTTCAGGAACAAGCAAATTTCACGTTGACTTGGAAAAAGAACTGGCTGATCTTCATGGAAAAGATGCAGCCTTGTTGTTCTCATCTTGCTTTGTAGCCAATGATTCCACCCTCTTCACTCTTGCTAAAATGCTGCCAGGTTGTGAGATCTACTCTGATTCTGGAAACCATGCCTCCATGATCCAGGGCATTCGAAACAGCAGGGTGCCGAAACACATCTTTCGCCATAACGATGTCAACCATCTTCGAGAGCTATTGAAGAAGTCTGATCCGTCTACCCCTAAAATTGTTGCATTTGAAACTGTGCATTCTATGGATGGTGCTGTCTGCCCTCTGGAAGAGCTGTGTGACGTGGCTCACGAGCACGGGGCAATCACTTTTGTGGATGAAGTGCACGCTGTGGGGCTGTATGGAGCTCGAGGTGGTGGGATAGGGGACCGCGATGGAGTCATGCACAAGATGGACATCATCTCTGGAACTCTTGGCAAGGCCTTTGGCTGTGTGGGTGGATACATCTCCAGTACAAGTGCTCTGATAGACACTGTCCGTTCATATGCTGCTGGTTTCATTTTCACTACATCCCTGCCACCCATGCTCCTGGCTGGTGCCCTCGAATCTGTCCGAACTCTGAAAAGTGCTGAGGGACAAATCTTGAGGCGCCAGCACCAACGCAATGTGAAGCTTATGAGACAGATGCTGATGGACGCAGGGCTTCCTGTAGTACACTGCCCAAGTCACATCATTCCAATAAGGGTTGCAGATGCTGCTAAAAATACAGAGATCTGTGACAAGCTGATGAGCCAGCACAGCATCTATGTCCAAGCCATCAACTACCCCACAGTTCCTCGTGGAGAAGAGCTGCTACGTATTGCTCCTACACCTCACCACACCCCTCAGATGATGAGTTATTTCCTTGAAAAGCTGCTGGCTACATGGAAGGATGTTGGGCTGGAGCTGAAGCCTCACTCATCAGCTGAATGCAACTTCTGCAGAAGACCTCTCCACTTCGAAGTGATGAGTGAAAGGGAAAGATCCTACTTCAGTGGCATGAGCAAACTAGTATCTGTCAGTGCATGAAAGTAATGGTGTTC-3′.

**Sequence of *HEPHL1* cDNA used for *in situ* hybridization (nt): (1,434 bp)**

5′-AAACACCTACAGATGGAATGTCCCAGAGCAATCAGGCCCTGGGAAAACAGATCCCAACTGTATCACTTGGGTTTACTATTCAACAGCAAATTTCGTCAAGGACACGTATAGCGGTCTGATTGGTCCCCTTGTAGTCTGCAGAAAAGGAGTTCTAGATGAAAATGGCCAGAGAAAAGACATTGATCGTAAATTTACTCTTCTGTTTATGGTGTTCGATGAAAACAAATCCTGGTATTTGGAAGAGAACATTGAAACATACCTTCACAAGAGTCCCGATGATTTTAATTCCACTAAGAACTTTGTAGAAGGCAACAGCAAGCATGCCATCAATGGGAAGATTTATAACAGTCTCCTGGGTCTAACCATGAATGAAGGGGATACGACAAACTGGTATTTGATAGGAACGGGTAATGAAGTAGATGTGCATACAGTCCATTTCCATGCACAGACCTTCATCTTCAAGACAGATAAAGACCACAGAGGAGATGTATATGACCTTTTCCCTGGGACTTTCCAAGCTGTTGAACTTGTAGCGGAAAACCCTGGAACGTGGCTTCTGCACTGCCACGTAGCTGACCACATACACGCTGGCATGGAAACGACCTACACCAACAATAAATCAGAGCTGGAAGCCCCTTCAGAAGGAGGACTGACGACCACCACAGCTTATGGTACAACTACCGCACACAACAGGACCACTGCAAAGGATGCTGACAGCCAAGGGGACAACGTCGGCTCTCCATGACAAAAGTGCACTTCTAGCAGATTCCCTGTAAATCCCCACTTTCCATAGAGCCATGCCGGGTTTCACCAAATGCTGCTGCTACAACTAGAGCAACACACACTTCAGCATTCCCATCCCATTGCCAGCACTCCTGAGAGCTGCAGGAAAGTGCTTCAGGAGCCACGTATCACCCATTGCCATTCCCACCCCACCACCACTCAGCTGTGCCCAAGGCACCACACCACTTGGCCAGCATTCTCATGGAGCAGACAGGATTGCACCCATGTGCTCTATGCAGCACCATGGAGTGCACACTGCTTTAGCATCCAGCTTCTGAAGGCACAGGTGTAAGTCTGGAAGTTCTTAAGGTTTCCCAGAGAGAAGCCAGCCTTACACCAGAGTCTGCATCAATGAATTCACCAGCATTTAACCGATTTTTCCCTTTTTTTGGTTCTCTTCACCCTGTTAAATGAGGAAGCTGTGCACTGGGGACAGATTTTTAAAGGCATTTAAGTAGCCACCAATATACTTCAGCACCTGATGAAGCTCAGCATCTGCAACAGGTTTGTTAATGCCTTTAAAAATCTAGCCCTGAGCAGCCTCAATGAAGTTATACTCTGTGATTGCTGCTTGTTGTGGTCCTGATTCTGCAACTCTTCTCTGCACTGAGCAGTGCTTTGTTCCCAAGCGTTAATTGGGCTGCTGATATC-3′

**Sequence of *PITX3* cDNA used for *in situ* hybridization (nt): (1,101 bp)**

5′- ATTCCTGCCCCATGGATTTCAACCTGCTGGCGGACGCGGAGGCTCGCAGCCCAGCCCTGTCCCTCTCAGACTCTGGCACCCCCCAGCACGAGCACAGCTGCAAGGGGCAGGACCATAGCGATACTGAGAAGTCCCAGCAGAACCAGACAGACGACTCCAACCCCGAGGACGGCTCTCTCAAGAAGAAGCAGCGGAGGCAGCGGACGCACTTCACCAGCCAGCAGCTCCAGGAGCTGGAAGCCACGTTCCAGAGGAACCGCTATCCTGACATGAGCACCAGGGAGGAGATTGCAGTCTGGACCAACCTGACGGAGGCACGAGTGCGGGTCTGGTTCAAAAATCGCAGGGCTAAGTGGAGGAAACGGGAGAGGAACCAGCAGGCCGAGCTCTGCAAGAACAGCTTTGGAGCCCAGTTCAATGGACTGATGCAGCCCTATGATGACGTGTATTCCAGCTATTCCTACAACAACTGGGCCACCAAAGGGCTTGCCACCAGCCCGCTCTCAGCTAAAAGCTTCCCATTCTTCAACTCCATGAACGTCAGCCCCCTCTCCTCCCAGCCCATGTTCTCCCCACCCAGCTCCATCGCCTCGATGACCATGCCTTCATCCATGGTCCCTTCTGCAGTGACTGGCGTCCCGGCCTCCAGCCTCAACAACCTGGGAAACATCAACAACCTGAACAGCCCAAGCCTCAACTCTGCCGTCTCATCCAGTGCCTGTCCTTACGCCTCCACAGCCAGCCCCTACATGTACAGGGACACGTGCAACTCCAGCCTGGCAAGTCTGAGGTTGAAGGCCAAGCAGCATGCCAACTTTACTTACCCGGCGGTGCAGACGGCAGCTTCCAACCTAAGCCTTGCCAATACGCCGTGGACAGGCCTGTATGAAGTGCTGCTCTCTGCTGACTGGGGACTTGACCTTAGCCTGACACAAGGAGACCTTTAGCCCTACAACAGTGTACATCCTGCAGAGCAAGAGTGGAAGCAAGAGAGAGCAAAAGAGCATGAGAAAGTGAGGGCACACAGCAAGCAGGCAGATGGACCTCGATGAAGATTTGTCTAACTATCGTATGGTGAATTTTGACTGTCTTTCCCCTCCC-3′.
